# The photoreaction process of a large-Stokes-shift red fluorescence protein-LSSmCherry1

**DOI:** 10.1101/2025.04.01.646287

**Authors:** Gaoshang Li, Jiajia Meng, Xiaolu Bai, Siteng Zhao, Yongnan Hu, Jin Dai, Yin Song, Xubiao Peng, Qing Zhao

## Abstract

Large Stokes shift red fluorescent proteins are highly valued in fluorescence imaging due to their considerable spectral separation and minimal self-absorption. However, a notable gap remains in our understanding of these proteins’ excited-state dynamics. Unlocking this knowledge could potentially drive significant advancements in cellular and molecular biology. In this study, we systematically examine the excited-state dynamics of LSSmCherry1, a large Stokes shift red fluorescent protein, under varying isotopic compositions and temperatures. Our aim is to elucidate its distinctive spectral properties. Through steady fluorescence spectral experiments at different temperatures, we demonstrate that the large Stokes shift is significantly reduced by approximately 80 nm at low temperatures compared to room temperature. Using transient fluorescence and absorption spectroscopy, we dissect the excited-state dynamics of LSSmCherry1 in neutral environments. We also investigate the kinetic isotopic effect by comparing spectra in water and heavy water. These analyses allow us to delineate plausible photocyclic pathways. Our results reveal that the excited-state dynamics of LSSmCherry1 follow a model similar to that observed in other fluorescent proteins, such as Green Fluorescent Protein (GFP). Notably, we demonstrate that the excited-state proton transfer (ESPT) process is the primary origin of the large Stokes shift in LSSmCherry1. This ESPT process occurs rapidly, within approximately 365 fs after excitation in the neutral environment. This study provides crucial insights into the mechanisms underlying large Stokes shift fluorescent proteins, potentially paving the way for the development of improved fluorescent probes for biological imaging.

**TOC Graphic:** Some journals require a graphical entry for the Table of Contents. This should be laid out “print ready” so that the sizing of the text is correct. Inside the tocentry environment, the font used is Helvetica 8 pt, as required by *Journal of the American Chemical Society*.

The surrounding frame is 9 cm by 3.5 cm, which is the maximum permitted for *Journal of the American Chemical Society* graphical table of content entries. The box will not resize if the content is too big: instead it will overflow the edge of the box.

This box and the associated title will always be printed on a separate page at the end of the document.

## Introduction

Large Stokes shift red fluorescent proteins hold great promise in the field of cell imaging due to their unique properties. These proteins exhibit a significant Stokes shift between their excitation and emission wavelengths, effectively reducing self-absorption effects and enhancing the signal-to-noise ratio in imaging applications.^1–5^ The pronounced Stokes shift and red fluorescence make these proteins particularly suitable for advanced imaging techniques, such as two-photon microscopy and multicolor imaging.^6–10^ Additionally, their use of the red emission spectrum offers advantages including reduced cellular autofluorescence and improved imaging depth, facilitating the observation of proteins and DNA within biological specimens.^11,12^

Understanding the mechanism and dynamics of the excited states in these proteins is crucial for optimizing their performance in imaging applications. Studies on GFP have revealed a three-state mechanism where excited-state proton transfer (ESPT) plays a pivotal role in generating vibrant green fluorescence.^13,14^ Investigations into large Stokes shift red fluorescent proteins such as mKeima, LSSmKate1, LSSmKate2, and Keima have also indicated ESPT involvement in their photophysics.^2,15–17^ Moreover, the Stokes shift of mPlum has been attributed to solvent reorganization,^18,19^ while the mechanism of Tag RFP675 relies on hydrogen bonding interactions.^20,21^

Innovative research led by Robert E. Campbell’s team has resulted in the development of novel large-Stokes-shift red fluorescent proteins, notably exemplified by LSSmCherry1, which is characterized by its impressive Stokes shift.^22,23^ LSSmCherry1, with its unique spectral properties, represents a significant advancement in the field of fluorescent proteins. However, the precise photophysical mechanisms underlying its large Stokes shift remain to be fully elucidated. The objective of the current study is to investigate the mechanism and dynamics of LSSmCherry1 in its excited state. To achieve this, we take a multifaceted approach, initially conducting thorough spectroscopic analyses that encompass investigations into isotopic effects and temperature dependencies using steady-state spectroscopy techniques. Subsequently, we utilize methodologies such as picosecond time-resolved fluorescence spectroscopy and femtosecond time-resolved transient absorption spectroscopy to dissect the evolution pathways of the excited states of LSSmCherry1. Through these comprehensive analyses, we aim to propose a detailed photocyclic process for LSSmCherry1, providing insights into its unique photophysical properties and paving the way for future rational design of improved fluorescent proteins.

## Results and discussion

### Influence of pH on Steady-State Spectral and Structural Characteristics

Figure 1A presents a schematic diagram of transient absorption spectroscopy. Currently, obtaining the crystalline structure of LSSmCherry1 remains challenging. In this study, a structural model was constructed through mutations based on the available structure of mCherry (PDB:2H5Q) using Rosetta software.^22,24,25^ Further details regarding this procedure are provided in the Methods section. Drawing upon insights from previous studies,^26,27^ amino acids S161 and E163 are identified as the likely critical residues responsible for generating large-Stokes-shift fluorescence.

**Figure 1:**
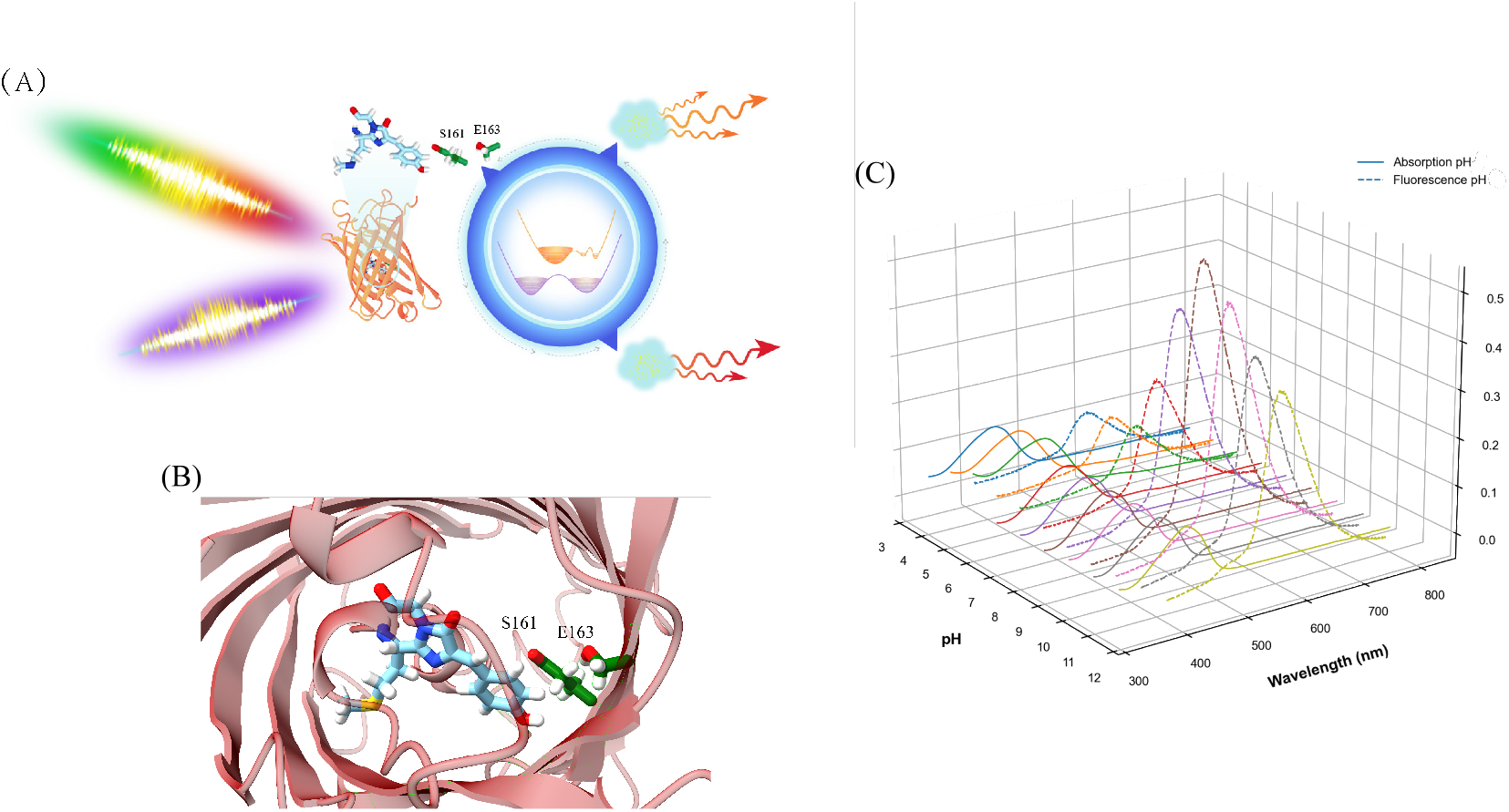
(A) Schematic representation of the transient absorption spectroscopy measurement process for LSSmCherry1. (B) Structural representation of the LSSmCherry1 chromophore within the -barrel. The image emphasizes the crucial residues involved in hydrogen bonding, which likely contribute to the protein’s large Stokes shift. (C) Three-dimensional plot illustrating the absorption and fluorescence spectra of LSSmCherry1 under various pH conditions.

Figure 1 illustrates the pH-dependent absorption and fluorescence spectra of LSSm-Cherry1, offering insights into its ground-state and excited-state electronic properties across a wide pH range (3-11). The absorption spectra (solid lines) reveal two distinct peaks: a predominant peak around 450-470 nm, indicative of the protonated state (A state), and a minor peak or shoulder near 570-580 nm, corresponding to the deprotonated state (B state). The relative intensities of these peaks vary with pH, reflecting the chromophore’s protonation equilibrium. The fluorescence spectra (dashed lines) exhibit a large Stokes shift, with emission maxima around 600-620 nm regardless of the excitation wavelength. This suggests that excited-state proton transfer (ESPT) plays a crucial role in the fluorescence mechanism of LSSmCherry1. The fluorescence intensity demonstrates a strong pH dependence, with maximal emission observed in neutral to slightly alkaline conditions (pH 7-8).

At low pH (e.g., pH 3), the absorption spectrum is dominated by the protonated A state, while the fluorescence intensity is relatively low. As pH increases, we observe a gradual increase in the B state absorption and a concomitant rise in fluorescence intensity. This pH-dependent behavior is consistent with other large Stokes shift fluorescent proteins, such as mKeima, LSSmKate1, and LSSmKate2.^15,17,28^ The spectral characteristics across the pH range suggest that LSSmCherry1 undergoes efficient ESPT, particularly in neutral to alkaline conditions. This process likely involves the conversion of the excited protonated state (A*) to an intermediate state (I*) and finally to the emissive deprotonated state (B*), accounting for the large Stokes shift observed.

According to the fluorescence spectra shown in Fig 1, the dynamics of excited-state proton transfer (ESPT) play a pivotal role in shaping the large-Stokes-shift characteristic of LSSm-Cherry1. To delve deeper into the underlying mechanism, we employed low-temperature experiments and isotopic substitution techniques. Specifically, we substituted hydrogen atoms with deuterium, which is known to affect the rate of ESPT. This modification in the fluorescence emission spectrum further highlighted the red emission band beyond 600 nm, indicative of anions generated via ESPT (Figure 2A). Subsequently, we examined the fluorescence spectrum at a low temperature of 80 K and pH 7 with excitation at 400 nm. As the temperature decreased, we observed a gradual intensification of the I band, along with a significant blue shift in the emission spectrum peak from 610 nm at 300 K to 530 nm at 80 K (Figure 2B). This substantial blue shift, which surpasses that observed in other Stokes-shifted fluorescent proteins, underscores the intricate relationship between temperature and ESPT dynamics.^29,30^ Furthermore, the temperature-dependent behavior of LSSmCherry1 also validated the existence of an energy barrier between the fluorescent states I* and B*. At the low temperature of 80 K, the system struggles to overcome the energy barrier on the excited-state energy surface from the intermediate state I* to the ionic state B*. Consequently, a higher portion of the intermediate state I*, which emits at shorter wavelengths, is observed. As the temperature increases to 300 K, which is sufficient for the system to easily overcome the barrier, the ionic state B*, characterized by longer emission wavelengths, becomes dominant.

**Figure 2:**
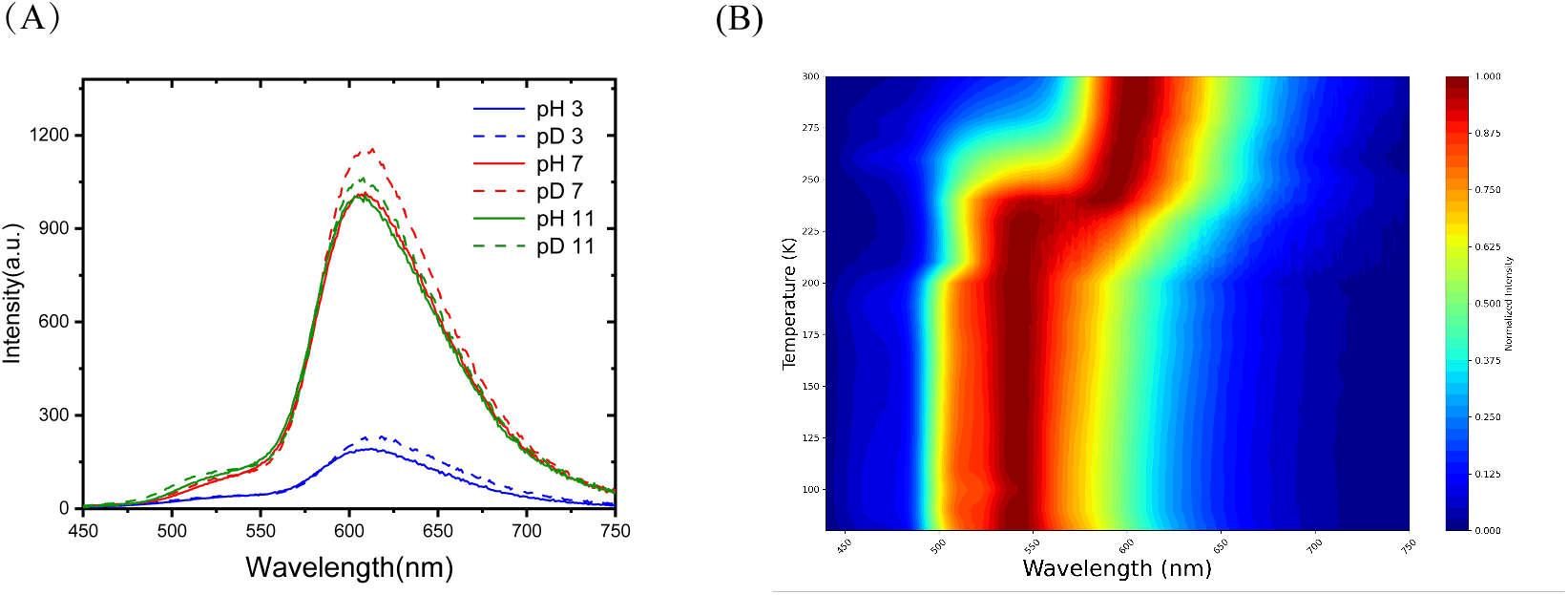
(A) Emission spectra are presented for pH 3 (blue), pH 7 (red), and pH 11 (green) in H_2_O (solid lines) and D_2_O (dashed lines). (B) Contour plot of normalized fluorescence spectra recorded at pH 7 across a temperature range from 80 K to 300 K, with measurements taken at 10 K intervals.

### Influence of pH and H/D Exchange on Fluorescence Dynamics

Our picosecond time-resolved fluorescence spectral measurements of LSSmCherry1 in both ordinary and heavy water buffers under neutral conditions (pH=7) reveal complex photophysical dynamics. Figure 3a presents raw data at representative wavelengths alongside curves fitted using a three-exponential model, demonstrating a pronounced wavelength dependence of fluorescence lifetime. The deconvolution of the fluorescence decay curves yields three distinct lifetime components: 1 (<100 ps), 2 (300-850 ps), and 3 (>1.0 ns). Notably, the amplitude of 1 decreases progressively as the emission wavelength shifts towards the red region. This trend corroborates our hypothesis from steady-state spectra that the A state is indeed unprotonated. Concurrently, the contribution of 3 increases significantly at longer wavelengths, suggesting a transition to a different emissive species.

**Figure 3:**
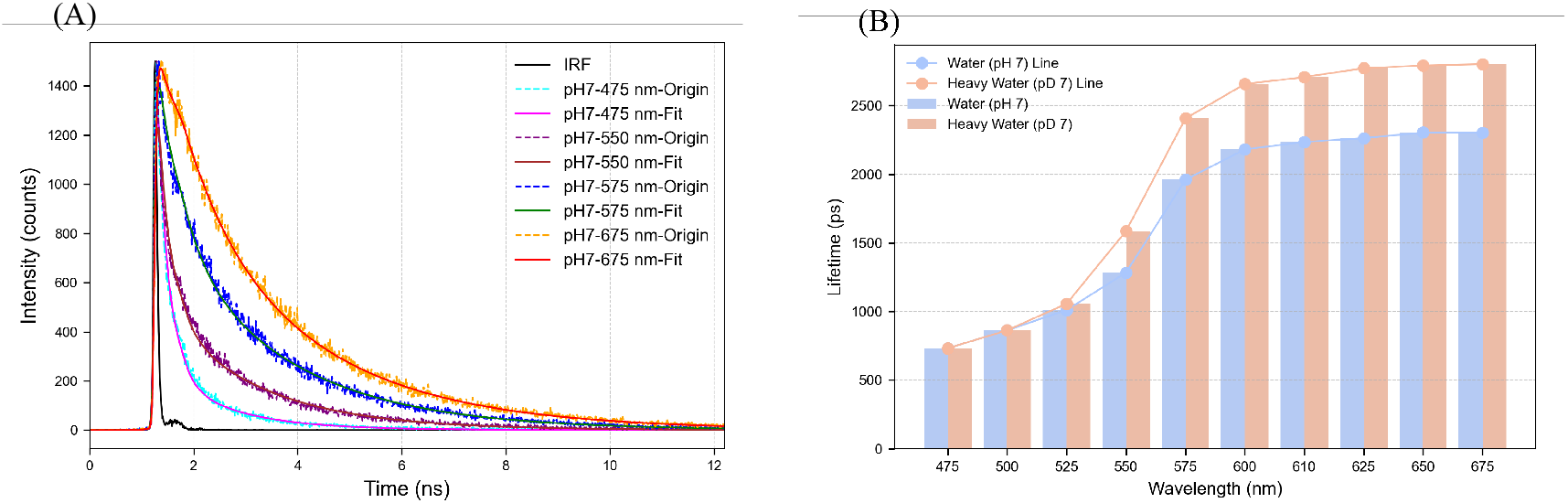
Time-resolved fluorescence analysis of LSSmCherry1 at pH/pD 7.0. (A) Fluorescence decay curves at various emission wavelengths (475 nm, 550 nm, 575 nm, and 675 nm) in pH 7 buffer. Solid lines represent fitted curves, while dotted lines show original data. The instrument response function (IRF) is shown in black. (B) Comparison of fluorescence lifetimes in H_2_O (pH 7) and D_2_O (pD 7) across emission wavelengths from 475 nm to 675 nm. Bar graph shows lifetimes, with connecting lines indicating trends.

Integrating these time-resolved results with steady-state fluorescence spectra allows us to attribute the short-lived component (1) to the excited-state emission of the neutral form A. This assignment is consistent with the low-intensity emission observed in the 480–540 nm range during steady-state experiments (Figure 3b). The intermediate (2) and long (3) components likely correspond to the I* and B* states, respectively, reflecting the progression of the excited-state proton transfer (ESPT) process. The isotope substitution experiment (Figure 3b) provides further insights into the proton transfer dynamics. The negligible effect on fluorescence lifetime at shorter wavelengths (450-500 nm) upon D2O substitution substantiates our assignment of the 450 nm band to the A* state emission. Conversely, the marked increase in lifetime at longer wavelengths (associated with anionic species) in D2O buffer strongly supports the ESPT mechanism as the primary conversion process in LSSmCherry1. This lifetime extension can be attributed to the kinetic isotope effect, where deuterium substitution alters the hydrogen bonding network around the chromophore, thereby slowing down the proton transfer rate. The multi-exponential decay and wavelength-dependent lifetimes observed here are indicative of a complex excited-state landscape. We propose a model where initial excitation populates the A* state, followed by rapid ESPT to form the I* state, which subsequently relaxes to the B* state. The relative populations of these states, as reflected in their amplitude contributions, evolve with emission wavelength, providing a spectral fingerprint of the ESPT process.

### Femtosecond Visible Absorption Spectroscopy to Detect Excited-State Dynamics

Transient absorption spectroscopy was employed to investigate the short-time dynamic processes in LSSmCherry1, offering insights beyond the resolution of time-correlated single-photon counting (TCSPC). We examined the excited-state dynamics of LSSmCherry1 in neutral environments using both water and heavy water. In Figures 5A: time-resolved spectra of LSSmCherry1 following excitation with a 400 nm laser pulse at pH 7 reveal distinct spectral features. A subtle ground-state bleaching (GSB) signal is observed below 480 nm, accompanied by an excited-state absorption (ESA) band from 500 to 580 nm and a stimulated emission (SE) band beyond 580 nm. At pH 7, both stimulated emission and excited-state absorption initially rise and subsequently decay. This evolution of initial enhancement followed by decay in the stimulated emission signal is attributed to excited-state proton transfer (ESPT), a phenomenon aligning with observations in other fluorescent proteins.^2,15–17^ Notably, within the first 100 ps, we observe an asynchronous evolution between the ESA and SE regions. This asynchrony likely results from the superposition of ESA with SE from the neutral excited state A*, consistent with our TCSPC results.

To quantitatively analyze the proton transfer process under neutral conditions, we employed global fitting to resolve the kinetic constants in water and heavy water.^31,32^ Using a sequential four-component model, we derived the decay-associated spectra (DAS) for H_2_O and D_2_O, as shown in Figure 4B and C. At pH 7, the waveform exhibits two rapidly rising components followed by a slower decaying component. An ultrafast process (365 fs) represents the transition from A* to I* via ESPT, followed by a subsequent proton transfer process (18 ps) leading to B* formation. A long-lived component (1089 ps) is attributed to the spectral decay of the excited anionic chromophore. To validate the proton transfer process, we performed global fitting in a heavy water environment at pH 7.

**Figure 4:**
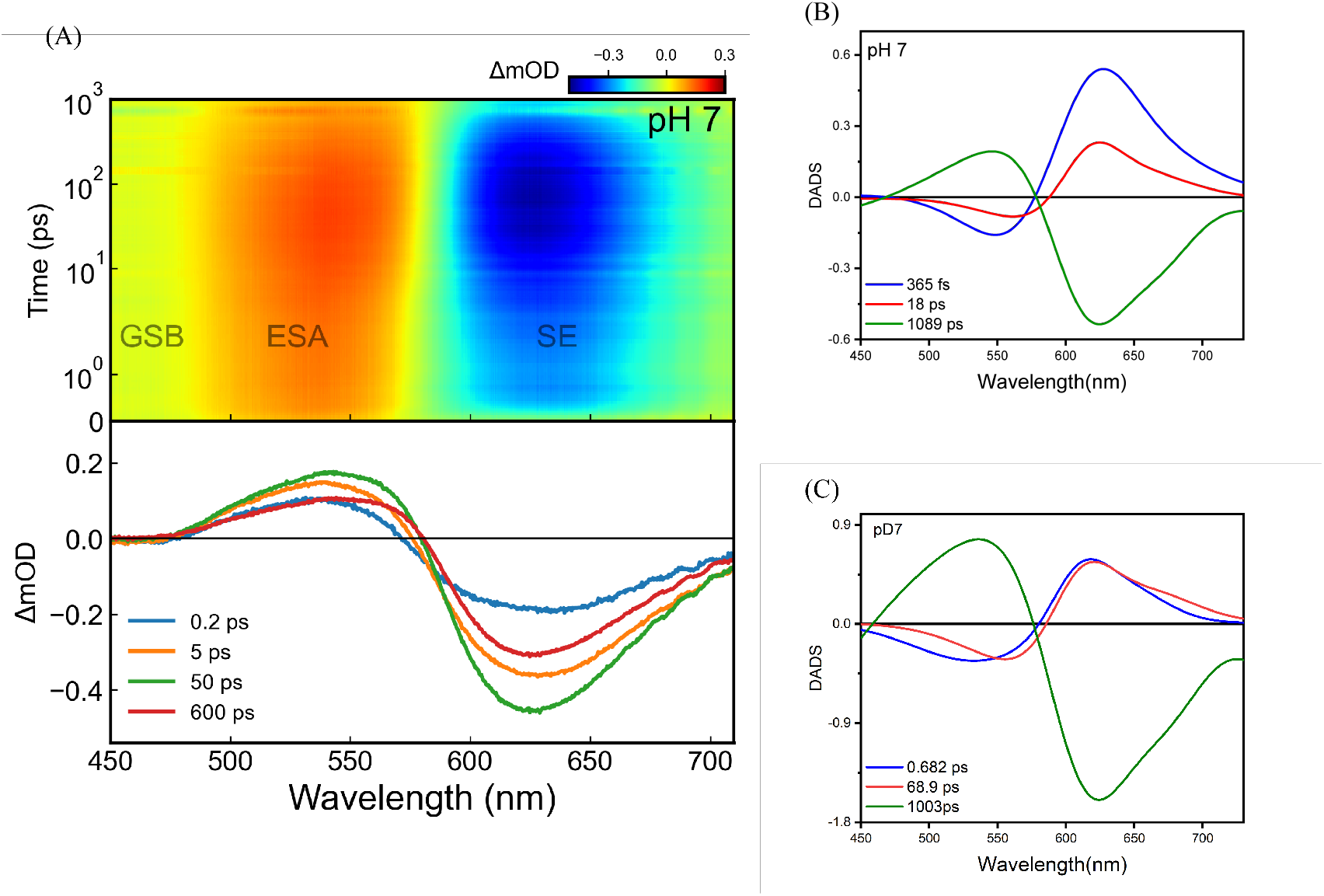
Transient absorption spectroscopy and global fitting analysis of LSSmCherry1 at pH 7. (A) Time-resolved absorption spectra in H_2_O following 400 nm excitation. Decay-associated spectra (DAS) obtained from global fitting analysis in (B) H_2_O and (C) D_2_O.

**Figure 5:**
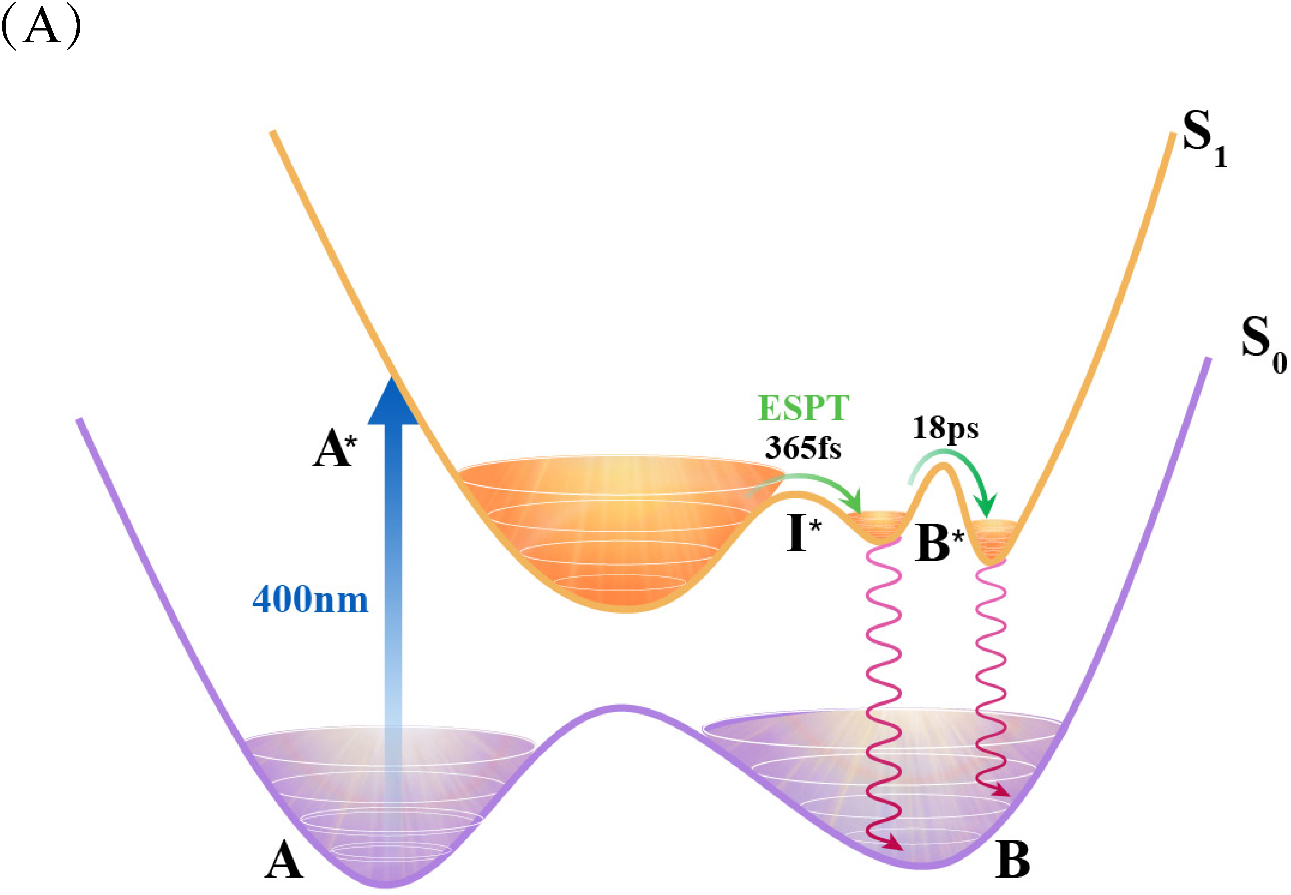
(A) Excited state proton transfer model of LSSmcherry1

The first two rising components (682 fs and 68.9 ps) showed significant kinetic isotope effects (KIE), confirming their association with proton transfer phenomena. The calculated KIE values (1.87 and 3.83 for the two components, respectively) further support the assignment of these processes to proton transfer events. Quantitatively, we estimate the ESPT rate constant to be approximately 2.74 *×* 10^12^ s^−1^ (per second) in H_2_O and 1.47 *×* 10^12^ s^−1^ in D_2_O at pH 7. These rates are comparable to those observed in other ESPT-exhibiting fluorescent proteins, such as GFP variants.

The complex dynamics observed can be related to the structural features of LSSm-Cherry1. The chromophore is likely situated in an environment that facilitates proton transfer, possibly involving a network of hydrogen bonds and strategically positioned amino acid residues. Future X-ray crystallographic studies could provide more detailed insights into this structure-function relationship. Compared to other red fluorescent proteins, LSSmCherry1 exhibits relatively fast ESPT kinetics, which may contribute to its large Stokes shift. This property sets LSSmCherry1 apart from traditional RFPs and makes it particularly useful for certain imaging applications, such as multicolor imaging or FRET-based studies.

Combining the results of all the above experiments, we give the conclusions as follows: 1. The steady absorption spectral experiments at different pH values reveal that the ground state of LSSmCherry1 predominantly exhibits a neutral chromophore with a peak absorption wavelength of approximately 447 nm, which is little influenced by the pH values. 2. The Gaussian decomposition of the steady fluorescent spectra indicates the existence of multiple fluorescent states, corresponding to the neutral state A*, intermediate state I* and ionic state B* on the excited-state energy surface, respectively. The percentages of the three states in fluorescence spectra vary as the pH value changes. The red fluorescence at 610 nm is primarily attributable to the formation of anionic chromophores in the excited state, which is further validated by the fluorescence spectral experiments in various temperatures. At low temperature, the peak of the fluorescent spectrum is blue-shifted by approximately 70 nm due to the high energy barrier between I* and B*. 3. At the room temperature, the solvent pH significantly influences fluorescence intensity, where the acid environment (pH=3) gives the weakest fluorescence. One possible reason is that the very low pH values can change the protonation status of the carboxylate group of E163, which further impacts proton transfer efficiency. 4.Through Time-Correlated Single Photon Counting (TCSPC) and transient absorption spectral experiments, we obtained kinetic parameters of the excited state evolution in LSSmCherry1. Isotope-dependent experiments further confirm that the Excited-State Proton Transfer (ESPT) process is essential for generating the large Stokes shift.

Based on a comprehensive analysis of all experimental phenomena, we propose the following photophysical process for LSSmCherry1 (refer to FIG 5): Following 400 nm pulse excitation, represented by the purple arrow, the neutral ground state species A is promoted to its excited state A* in the S1 electronic manifold. Subsequent to excitation, A* predominantly undergoes ESPT (Excited State Proton Transfer), rapidly forming an intermediate state I* with a time constant of 365 fs. This ultrafast proton transfer is a critical step in the photophysical cascade. The I* state serves as a pivotal juncture, branching into two competing pathways: it can either emit fluorescence, likely corresponding to the 600 nm emission band, or further evolve into the anionic excited state B*. The latter transformation occurs over a longer timescale of 18 ps, indicating a more gradual structural rearrangement. The anionic B* state, once formed, radiatively decays to its ground state B, presumably giving rise to the red-shifted fluorescence emission at approximately 640 nm. While not explicitly shown in the figure, it’s plausible that a minor fraction of A* may directly relax to the ground state, potentially accounting for the weak fluorescence observed around 520 nm. We further analyze the possible ESPT pathway according to the computationally optimized structures using Rosetta.^22,24,25^ From Figure 1B, we can see that the residues S161 and E163 in the LSSmCherry1 are deemed close to the chromophore, which further validates the proposed mechanism in reference^23^ that the hydroxyl group of the sidechain S161 and the carboxyl group of the sidechain in E163 play crucial roles in facilitating the deprotonation and reprotonation processes.

## Conclusion

In summary, our investigation of the fluorescent protein LSSmCherry1 elucidated that its significantly LargeStokesshift primarily arises from the ESPT process, with rapid initial proton transfer occurring at 365 fs, followed by a slower transition to the anionic state at 18 ps. LSSmCherry1’s mechanism resembles that of other fluorescent proteins with significant Stokes shifts, such as Keima or LSSmOrange. Our thorough analyses demonstrated the considerable influence of temperature, variations in pH, and hydrogen/deuterium isotopes on the LSSmCherry1 Stokes shift. We’ve proposed a detailed photophysical model involving multiple excited states (A*, I*, and B*) and their associated transitions. However, the potential influence of ground-state heterogeneity on dynamics could not be ruled out, which would require the support of subsequent crystal structures. The observed weak emission around 520 nm might indicate a minor direct relaxation pathway from A* to the ground state, warranting further investigation. Overall, our study provides profound insights into the properties and photophysical processes of this large-Stokes-shift red fluorescent protein, offering valuable guidance for the rational design of improved variants with tailored spectral characteristics.

## Experimental methods and means

### Structure Modeling

For each mutant, we utilized Rosetta software^22,24,25^to conduct mutation modeling starting from the nearest structure (PDB:2H5Q). Following the addition of constraints on the C-*α* coordinates, we executed four rounds of Repack-MinMover cycle iterations. Thirty structures were concurrently generated, and the structure exhibiting the lowest Rosetta energy (r.e.u.) was designated as the final structure, determined through evaluation utilizing the Rosetta all-atom scoring function.^26^

### Protein Expression, Purification and Deuteration

The LSSmCherry1 gene fragment was synthesized using whole gene synthesis and subsequently inserted into the *pet-30a* vector. The vector was then transformed into *E. coli* cells for expression and purification. The specific procedure is outlined as follows: induction of logarithmic growth at 37^°^C in a medium containing kanamycin, followed by the addition of 0.5 mM IPTG to induce expression for four hours. Subsequently, the cells were centrifuged at 4000 rpm for 15 minutes to collect the pellet, which was then resuspended. Cell disruption was achieved through sonication after rapid freezing in liquid nitrogen. Protein purification was conducted using Ni-NTA agarose (Qiagen), and the elution buffer was exchanged with Tris-Hcl buffer (25 mM Tris, 150 mM NaCl) using desalting columns (GE Healthcare). The purified proteins were temporarily stored at 4^°^C or flash-frozen and stored at −80^°^C for long-term storage. Deuteration of exchangeable protons in the protein was carried out by dissolving the protein in a 25 mM Tris, 150 mM NaCl buffer in D_2_O at pH 8, concentrating to a minimum volume, and repeating the procedure eight times. It is noteworthy that the pH conditions were not adjusted for the isotope effect.

### Steady-state absorption, fluorescence spectroscopy, and low-temperature fluorescence spectrum measurements

To explore the pH dependency of spectra, we utilized buffers covering a pH range from 3 to 11. A portion of the purified protein was adequately diluted in the respective buffer solution. Spectra were recorded at room temperature (25°C) in Tris-Hcl(25 mM Tris, 150 mM NaCl) buffer. For the steady-state absorption spectrum, we prepared two parallel sets of samples for each pH value and measured their spectra. The actual final pH of each sample was determined using a microelectrode. Simultaneous measurement of steady-state absorption spectra and fluorescence spectra was performed using a Dual-FL spectrometer by HORIBA.

Low temperature fluorescence spectrum were measured using an Edinburgh Instruments time-correlated single photon counting (TCSPC) system(FLS1000). The protein concentration employed in the aforementioned experiment was 0.2 mg/mL.

### Fluorescence lifetime measurement

Fluorescence lifetime data were acquired employing a commercial time-correlated single photon counting (TCSPC) system (FLS980) featuring a 400 nm pulsed laser diode head for excitation. An emission filter, centered at 400 nm, was applied to eliminate excitation and scattered photons. All measurements were conducted in cuvettes with an optical path length of 1 cm at an optical density of approximately 0.1. The goodness of fit was assessed based on fit parameters, specifically the reduced chi-square (*χ*^2^), with values consistently below 1.2 considered indicative of high-quality fits.

### Femtosecond Time-resolved Absorption Measurement and Global Fit

The experimental setup closely followed the previously described configuration.^33^ Utilizing a Ti:Sapphire Regen amplifier, 800 nm laser pulses with a duration of 140 fs served as the primary light source to illuminate both the TOPAS-C and sapphire crystals. This laser pulse was divided into two beams, designated as the pump pulse and probe pulse. White light detection was achieved using CaF_2_. The pump light, generated through an optical parametric amplifier (TOPAS-C), reached 200 nJ at 400 nm. The polarization of the pump pulse was set to the “magic angle” (54.7°) relative to the horizontally polarized probe pulse. Both pump and probe pulses were efficiently focused into a 1 mm thick flow cell, where the sample solution circulated to ensure continuous replenishment with fresh solution between successive laser irradiations. The experiment utilized a protein concentration of 1.5 mg/mL in a buffer solution containing 25 mM Tris and 150 mM NaCl. For the transient absorption spectrum, we performed independent measurements for each sample, and finally took the average of the two results. The signals measured two times showed no significant deviation, indicating the stability of the sample during the measurement process and ensuring the reliability of the transient absorption spectrum signal. Data analysis and global fitting were executed using the KiMoPack package in Python.^31,34^

## Acknowledgement

Y.S. acknowledges support from the NSFC (No. 62105030). We acknowledge the Protein Preparation and Characterization Core Facility of Tsinghua University Branch of China National Center for Protein Sciences Beijing for providing the facility support. We thank the Biological and Medical Engineering Core Facilities of Beijing Institute of Technology for supporting experimental equipments.

## Supporting Information Available

The Supporting Information is available free of charge at https://pubs.acs.org.

Steady-state absorption and fluorescence spectra under various pH conditions (Figures S1 and S2); Gaussian decomposition analysis of fluorescence spectra (Figure S3); femtosecond transient absorption spectra and global fitting results (Figures S4 and S5); relative amplitudes from TCSPC data (Table 1); time constants and relative amplitudes from global fitting analysis (Table 2).

## Notes

### Competing Interest Statement

The authors have declared no competing interest.

