## Supplementary material for "The photoreaction process of a large-Stokes-shift red fluorescence protein-LSSmCherry1": SI file

### Content

• **Figure S1:** Steady-state absorption spectra under various pH conditions.

• **Table 1:** Relative amplitudes (Ai) were obtained from the average lifetime of the single-photon counting (TCSPC) spectroscopy data of LSSmCherry1 at pH 3, 7, and 11.

• **Figure S2:** Steady-state fluorescence spectra under various pH conditions.

• **Figure S3:** Gaussian decomposition analysis of steady-state fluorescence spectra at different pH levels.

• **Figure S4:** Femtosecond transient absorption spectra.

• **Figure S5:** Global fitting.

• **Table 2:** Time constants and their relative amplitudes (Ai) were obtained from the global fitting analysis of the time-resolved absorption data of LSSmCherry1 at pH 3, 7, and 11.

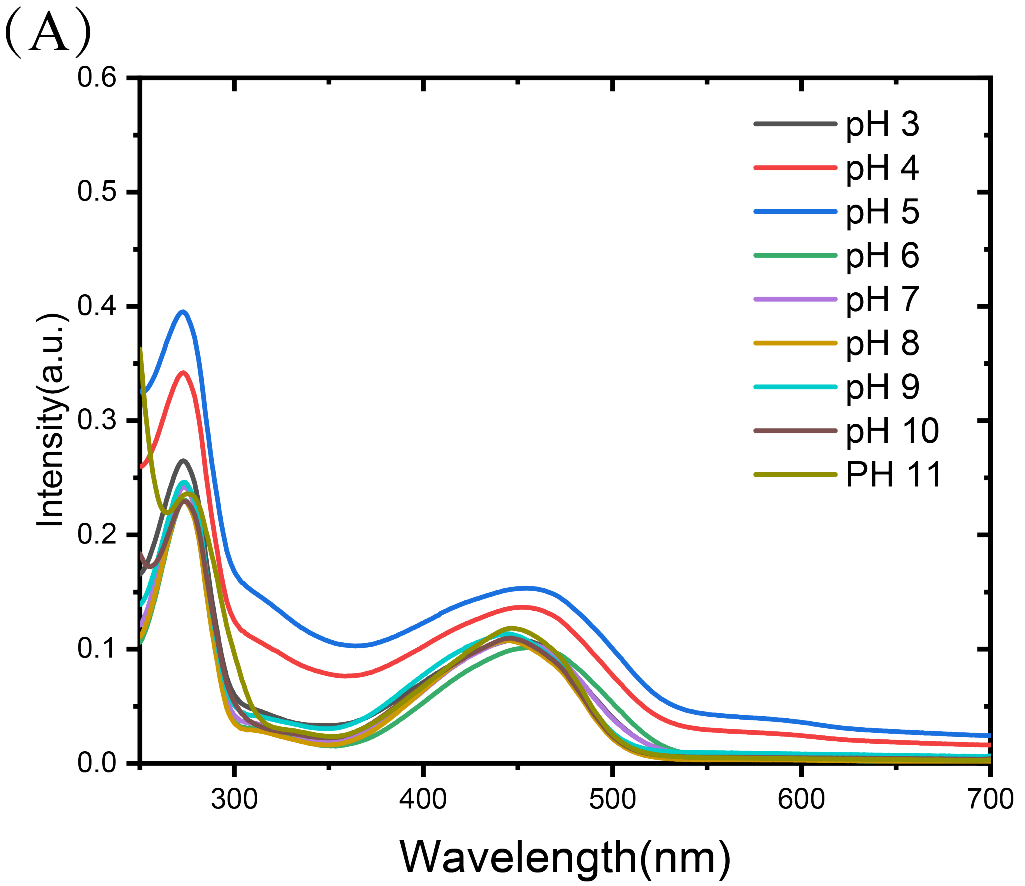


Figure 1: Steady-state absorption spectra under various pH conditions.

Table 1: Relative amplitudes ($A_i$) obtained from the average lifetime of the TCSPC spectroscopy data of LSSmCherry1 with fixed lifetimes, at pH 3, 7, and 11.

| **pH** | $\lambda$**(nm)** | **A1 (%)-53.36 ps** | **A2 (%)-448.97 ps** | **A3 (%)-1618.90 ps** |
|---|---|---|---|---|
| | 475 | 19.5 | 25.16 | 55.33 |
| **3** | 575 | 16.60 | 43.10 | 40.30 |
| | 610 | 5.73 | 37.89 | 56.38 |
| | | **A1 (%)-50.95 ps** | **A2 (%)-586.73 ps** | **A3 (%)-2120.37 ps** |
| | 475 | 24.59 | 55.13 | 20.28 |
| **7** | 575 | 3.28 | 12.25 | 84.47 |
| | 610 | 1.26 | 2.51 | 96.23 |
| | | **A1 (%)-48.08 ps** | **A2 (%)-579.10 ps** | **A3 (%)-2138.89 ps** |
| | 475 | 33.53 | 45.37 | 21.10 |
| **11** | 575 | 3.60 | 16.89 | 79.51 |
| | 610 | 1.34 | 6.00 | 92.86 |

*Note:* The fixed lifetimes used in fitting were taken from the average lifetimes across various wavelengths in Table 1 for each component at every pH level.

(A)

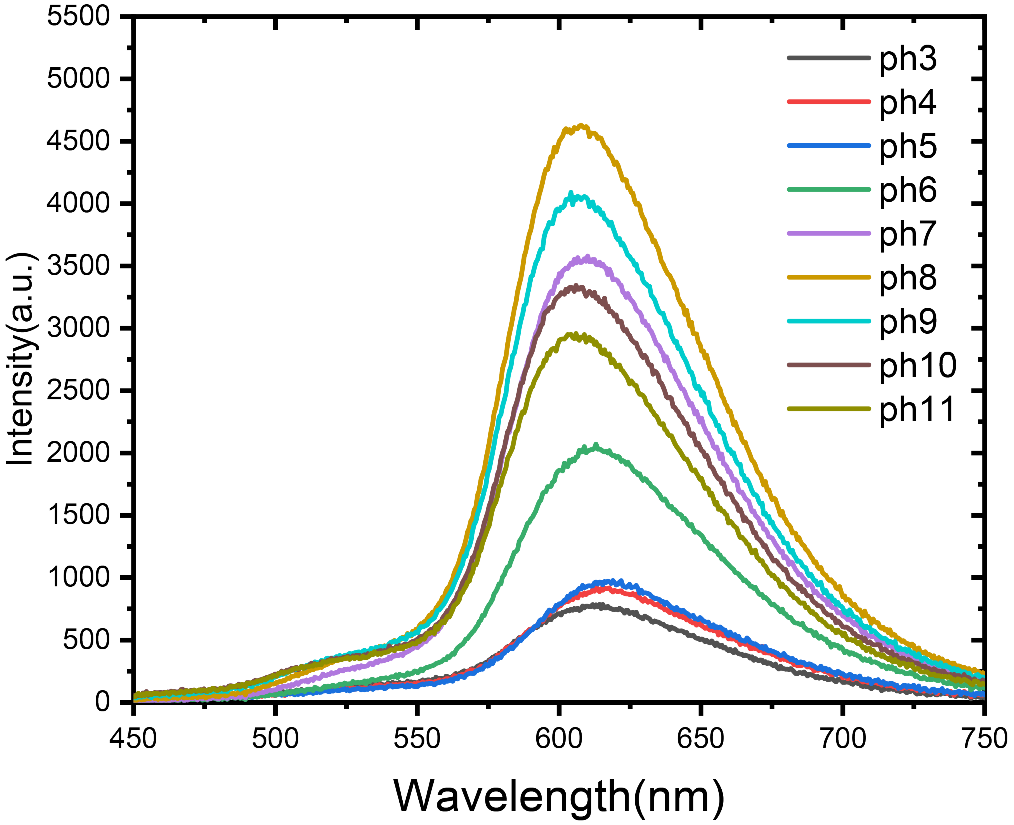


Figure 2: Steady-state fluorescence spectra under various pH conditions.

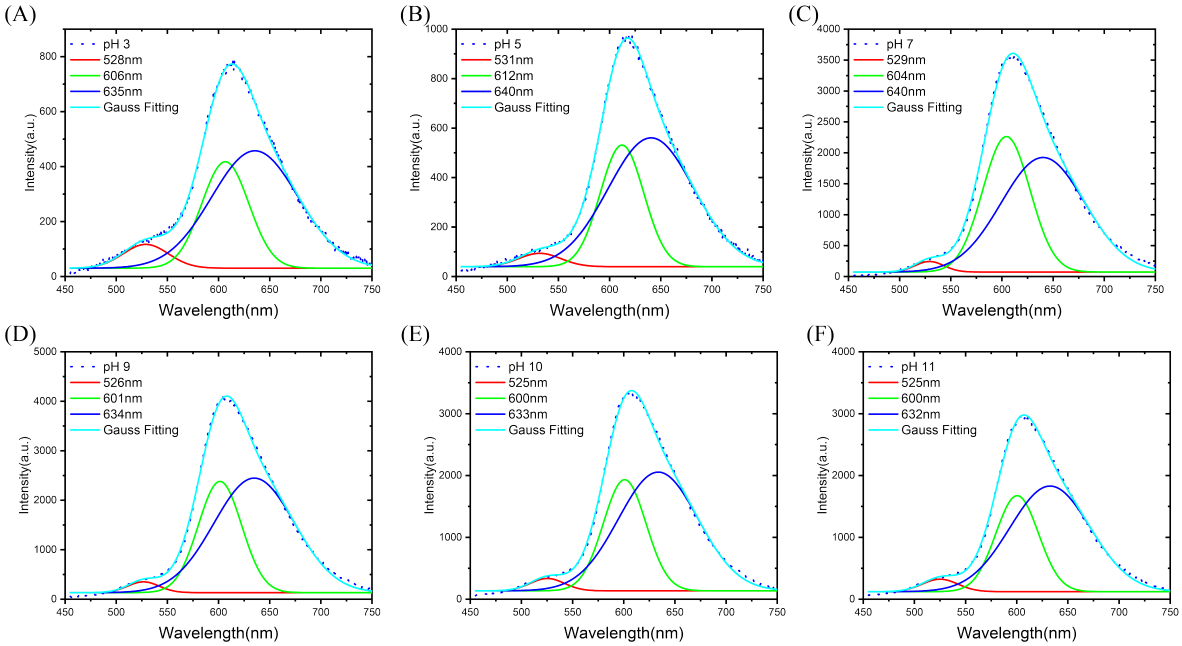


Figure 3: Gaussian decomposition analysis of steady-state fluorescence spectra at different pH levels. (A) Gaussian decomposition of the steady-state fluorescence spectrum at pH 3, (B) pH 5, (C) pH 7, (D) pH 9, (E) pH 10 and (F) pH 11.

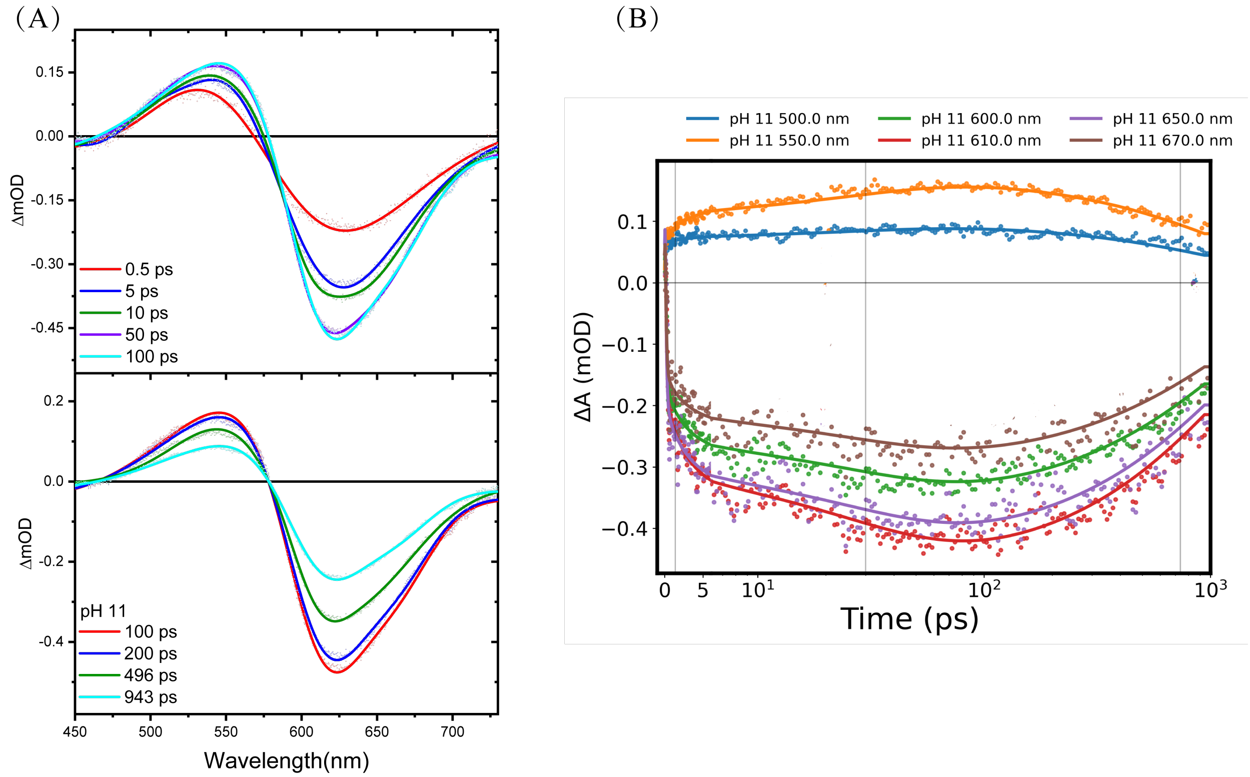


Figure 4: Femtosecond transient absorption spectra. (A) Femtosecond transient absorption spectra for LSSmCherry1 were recorded at pH 11 following excitation at 400 nm. (B) Kinetic traces at representative wavelengths at pH 11 is depicted, with symbols denoting measured data points and thick lines representing global fits.

Table 2: Time constants and their relative amplitudes ($A_i$) obtained from the global fitting analysis of the time-resolved absorption data of LSSmCherry1 at pH 3, 7, and 11.

| | **Wavelength (nm)** | **A1 (5.52 ps)** | **A2 (98.8 ps)** | **A3 (516 ps)** |
|---|---|---|---|---|
| | 500 | 0.140 | -0.116 | -0.744 |
| | 550 | 0.090 | -0.170 | -0.740 |
| | 600 | -0.082 | 0.153 | 0.765 |
| pH=3 | 610 | -0.147 | 0.022 | 0.831 |
| | 650 | -0.069 | 0.008 | 0.923 |
| | 670 | -0.020 | 0.030 | 0.950 |
| | | **A1 (0.365 ps)** | **A2 (18 ps)** | **A3 (1090 ps)** |
| | 500 | 0.448 | 0.099 | -0.453 |
| | 550 | 0.199 | 0.205 | -0.596 |
| | 600 | -0.231 | -0.192 | 0.577 |
| pH=7 | 610 | -0.334 | -0.193 | 0.473 |
| | 650 | -0.122 | -0.227 | 0.651 |
| | 670 | -0.080 | -0.230 | 0.690 |
| | | **A1 (2 ps)** | **A2 (30.8 ps)** | **A3 (1280 ps)** |
| | 500 | 0.121 | 0.207 | -0.672 |
| | 550 | 0.105 | 0.236 | -0.659 |
| | 600 | -0.145 | -0.199 | 0.656 |
| pH=11 | 610 | -0.135 | -0.230 | 0.635 |
| | 650 | -0.117 | -0.199 | 0.684 |
| | 670 | -0.113 | -0.195 | 0.692 |

*Note:* The amplitudes were normalized by dividing the sum of the amplitudes (A1 + A2 + A3) at each wavelength.
